# Direct Evidence for Dendritic Spine Compensation and Regeneration in Alzheimer’s Disease Models

**DOI:** 10.1101/2025.09.01.673437

**Authors:** Nishita Bhembre, Zoran Boskovic, Jessica Willshaw, Tim Castello-Waldow, Calum Bonthron, Annalisa Paolino, Patricio Opazo

## Abstract

Dendritic spine loss in Alzheimer’s disease (AD) strongly correlates with cognitive decline, whereas spine preservation is associated to cognitive resilience. Yet, whether and how neurons compensate for spine loss in AD remains largely unknown. Using a chromophore-assisted light inactivation approach (CALI), we developed a tool to selectively eliminate dendritic spines to model this key feature of AD. Using in vivo and in vitro two-photon imaging, we discovered that the artificial elimination of spines triggers a two-stage compensatory response: rapid enlargement of remaining spines followed by delayed spine regeneration. Remarkably, similar structural plasticity was observed across multiple β-amyloid-driven models of synapse loss, including the APP/PS1 mouse and following intracortical delivery of oligomeric β-amyloid. Mechanistically, compensatory spine enlargement required NMDA receptor activation and de novo protein synthesis. These findings suggest that neurons retain an intrinsic capacity to reverse early synaptic loss in AD, potentially contributing to cognitive resilience.

## INTRODUCTION

Dendritic spine integrity is a key determinant of cognitive function in Alzheimer’s disease (AD). Loss of dendritic spines tightly correlates with cognitive deficits in AD patients and animal models [10, 34, 40]. Conversely, the preservation of dendritic spines is associated with cognitive resilience — the ability to maintain normal cognition despite significant amyloid and tau pathology [2, 7, 16, 44]. Notably, restoring dendritic spine density in AD animal models is sufficient to rescue cognitive function[32]. These findings underscore the need to elucidate the compensatory and repair mechanisms that counteract spine loss in AD [5, 26].

Although the mechanisms underlying dendritic spine loss are increasingly well understood, whether and how neurons compensate for spine loss remains largely unknown [5]. Because early synapse loss in AD is gradual and localized to dendrites in close proximity to plaques [6, 19, 36], it is unlikely to engage classical homeostatic plasticity mechanisms like synaptic upscaling [41], which require sustained and widespread reductions in neuronal firing. Instead, local forms of spine compensation in the vicinity of lost spines are more likely to be implemented in the early stages of AD.

Snapshots on fixed-preparations from postmortem AD subjects, as well animal models of AD, have shown that dendritic spine loss co-occurs with an enlargement of the neighbouring spines [4, 5, 10, 33, 35, 43]. Although this may correspond to a compensatory adaptation to preserve the excitatory drive of the affected dendrite [5, 10], this has typically been attributed to the preferential loss of small, vulnerable spines [4]. In this view, simply increasing the proportion of large spines could account for the population-level increase in average spine size. To distinguish between these two possibilities, it is crucial to perform longitudinal live imaging experiments to investigate whether the loss of dendritic spines leads to the emergence of compensation over time at the individual spine level [5].

Another key challenge in studying the mechanisms of spine compensation in AD lies in the inability to predict which dendritic spines will be eliminated, and consequently, which ones will undergo compensation. To overcome this limitation, in this study we developed an optogenetic tool for the artificial elimination of dendritic spine with high spatiotemporal control and identified a two-stage compensatory response: rapid enlargement of remaining spines followed by delayed spine regeneration. Strikingly, we observed similar structural plasticity across multiple *in-vitro* and *in-vivo* models of β-amyloid-induced synapse loss. These findings suggest that neurons retain an intrinsic capacity to reverse early synaptic loss, potentially contributing to cognitive resilience in AD.

## RESULTS

### Optogenetic spine elimination triggers a two-stage compensatory response

Because dendritic spine loss in AD occurs stochastically, it is not possible to predict which synapses will be lost or compensated. To achieve spatiotemporal control over spine loss, we developed an optical approach for the targeted elimination of spines by disrupting the actin cytoskeleton using chromophore-assisted light inactivation (CALI) [9]. Using biolistic transfection in organotypic slices cultures, we overexpressed DrebrinA, an actin-binding protein which is highly enriched at spines [13], fused to the genetically-encoded photosensitizer KillerRed, along with the structural marker EGFP. Following a two-photon baseline imaging session, we locally inactivated Drebrin::KillerRed using epifluorescent green light and reimaged the same dendritic region 24 hours and 1 week later. As depicted in Fig. 1a.b we found that Drebrin inactivation led to the artificial elimination of dendritic spines 24 hours later compared to no-light controls (Drebrin::KillerRed overexpression without light inactivation). Using post hoc immunostaining of endogenous PSD95, we found that spine elimination had no effect on the density of shaft PSD95 puncta suggesting that spine synapses did not relocate onto the shaft after spine elimination (Suppl. Fig 1a-c). To investigate whether artificial spine elimination led to the enlargement of the surviving dendritic spines, we measured the size of individual spines before and after Drebrin::KillerRed inactivation by measuring their integrated brightness (see Methods). Remarkably, we found that spine elimination triggered the enlargement of the remaining spines at the individual spine level (Fig. 1c.d). By plotting the extent of spine enlargement as a function of its initial size, we found that smaller spines were preferentially enlarged (Fig. 1e and Suppl. Fig. 2a), in agreement with their plastic nature [8, 21]. Importantly, to examine whether spine enlargement was paralleled by functional changes, we artificially eliminated spines in neurons expressing Drebrin::KillerRed together with the genetically-encoded calcium indicator GCaMP6s. As shown in Fig. 1h.j, Drebrin inactivation led to a robust enhancement of spontaneously evoked calcium transients in individual spines 24 hours post-inactivation.

**Figure 1.**
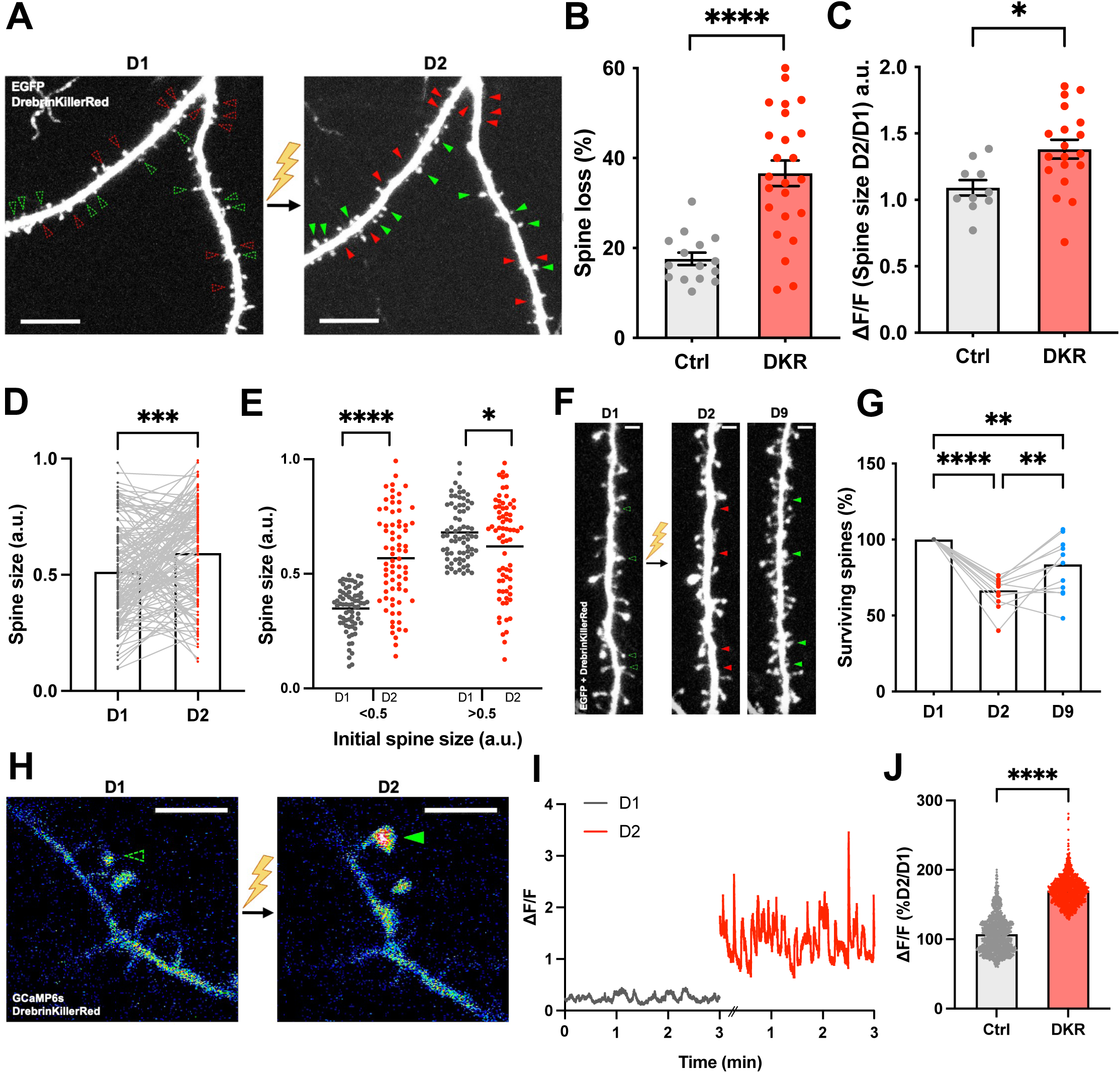
Artificial dendritic spine loss and compensation using a chromophore-assisted light inactivation (CALI) approach. a) Two-photon images of hippocampal neuron in organotypic slice cultures overexpressing GFP and Drebrin::KillerRed before and after the optical inactivation of Drebrin. Note that Drebrin inactivation leads to spine loss (red arrowheads) and spine compensation (green arrowheads) 24 hours post-inactivation (D2) as compared to baseline imaging (green/red hollow arrowheads). Scale bars 10 µm. b) Bar graph showing a significant loss of dendritic spines 24hours post-Drebrin inactivation Spine loss in controls (Ctrl)= 17.6± 1.373%, n=15 neurons; spine loss after Drebrin inactivation = 36.6± 2.86%. n=24 neurons ****p <0.0001 unpaired t-test. Data points in bar graph corresponds to neurons. All data are ± SEM. c) Bar graph showing a significant increase in spine size following Drebrin inactivation. Relative size increase between Day 2 and Day1 (D2/D1) in control (Ctrl) = 1.091±0.06 (n=10 neurons) and following Drebrin inactivation (DKR)= 1.381±0.07 (n=19 neurons). *p<0.05, unpaired Mann-Whitney t-test. Data points in bar graph corresponds to neurons. All data are ± SEM d) Bar graphs showing the increase in spine size at the individual spine level following Drebrin inactivation (Day1 =0.5133±0.17; Day2=0.5939±0.017). ***p <0.001 paired t-test (n=143 spines). e) Graph showing that smaller dendritic spines (brightness ratio < 0.5) are preferentially enlarged whereas large spines (brightness ratio >0.5) shrink after 24 hours. Small spines: Day1= 0.348 ± 0.011, Day2= 0.568±0.02; ****p <0.0001, n=72. Large spines: Day1=0.680±0.014, Day2=0.620±0.024; *p <0.05, n= 71 spines. f) Two-photon images of hippocampal neurons showing the regrowth of dendritic spines one week after Drebrin::KillerRed inactivation. Note that Drebrin inactivation induces spine loss (red arrowheads) and a compensatory regrowth (green arrowheads) of lost dendritic spines (green hollow arrowheads). Scale bars 2 µm. g) Bar graph representing the regrowth of dendritic spines following 1 week (Day9) post Drebrin inactivation when normalized to Day1 (Day 2 = 66.59±2.804 and Day 9= 83.69±5.13%, Day2/Day1 ****p<0.0001, Day9/Day1 **p<0.01, Day9/Day2 **p<0.01, one-way ANOVA, n=13 neurons. h) Representative dendritic region of a neuron overexpressing the genetically-encoded calcium indicator GCaMP6s and Drebrin-KillerRed. The GCaMP6s fluorescence signal in dendritic spines (green arrow) increased following Drebrin-KillerRed inactivation. i) GCaMP6s trace in the spine indicated with the green arrow in H) before and after Drebrin-KillerRed inactivation. j) Bar graph showing a significant increase in GCaMP6s signal in individual dendritic spines 24hours post Drebrin inactivation. Control=107.1±0.738%, DKR=170±0.552%. ****p<0.001 unpaired t-test (n=1270 traces)

To assess whether artificial spine elimination triggers spinogenesis as an additional form of structural compensation, we also quantified spine formation rates at 24 hours and 1 week following Drebrin inactivation. While no increase in spine formation was detected 24 hours after inactivation (Suppl. Fig. 2b), a substantial regeneration of dendritic spines became evident after 1 week (Fig. 1f.g).

### In-vivo validation of optogenetic dendritic spine elimination and compensation

Finally, we evaluated whether artificial spine elimination induces structural compensation *in vivo*. To that end, we delivered the optogenetic tool and the structural marker GFP in the cortex via *in-utero* electroporation at E15 and implanted a cranial window postnatally to allow for two-photon imaging and Drebrin inactivation (Fig. 2a). Following two consecutive baseline imaging sessions (Day 1 and Day 2), we locally inactivated Drebrin::KillerRed using epifluorescent green light and reimaged the same dendritic region 24 hours and 1 week later using *in-vivo* two-photon imaging. Although artificial spine elimination was less effective *in-vivo* than *in-vitro*—likely due to limited brain penetration of one-photon light— dendrites with >10% spine loss showed enlargement of surviving spines at 24 hours (Fig. 2e.f) and increased spine formation after one week (Fig. 2d).

**Figure 2.**
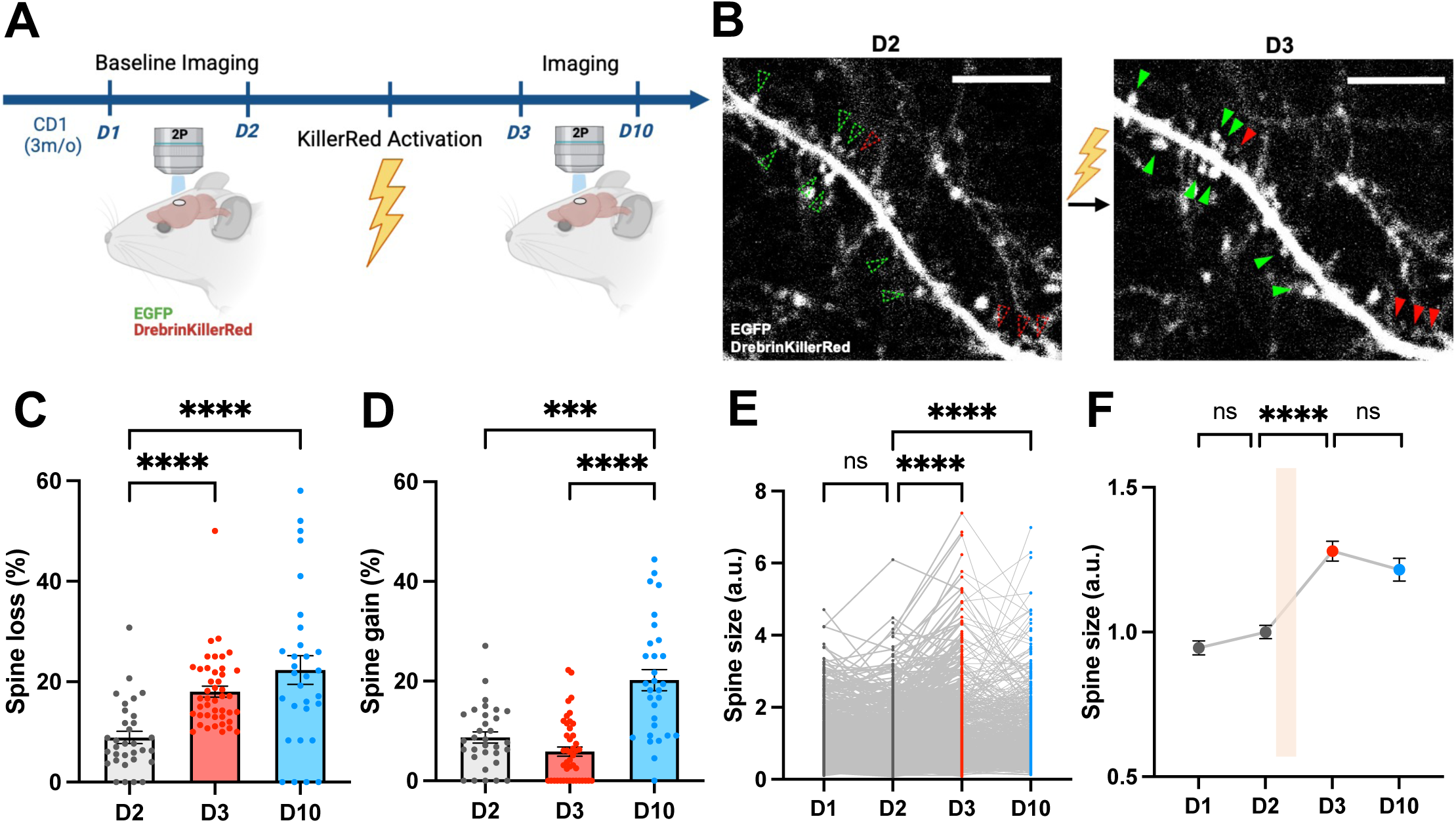
In-vivo validation of optogenetic dendritic spine elimination and compensation. a) Schematic displaying the experimental workflow for *in-vivo* two-photon imaging and inactivation of Drebrin in wild type CD1 mouse previously *in-utero* electroporated with Drebrin::KillerRed and EGFP. b) *In-vivo* two photon images of cortical neurons labelled with EGFP at baseline (Day 2) and 24 hours post-Drebrin inactivation (D3). Dendritic spine loss and compensation is indicated by red and green arrowheads, respectively, as compared to baseline (green/red hollow arrowheads). Scale bar 20 µm. c) Bar graph showing a significant loss of dendritic spines 24 hours (Day3) and 1 week (Day10) following Drebrin inactivation in dendrites with >10% spine loss (45/123 dendrites). Spine loss Day2 (baseline)=8.896±1.206%, n=32; Day3=18.03%± 1.091, n=44; Day10= (22.34±2.84%), n=30. Day2 vs Day3 ****p < 0.0001, Day2 vs Day10 ****p < 0.0001, Day3 vs Day10 non significant. Kruskal-Wallis test. d) Bar graph showing a significant increase in spine gain 1 week (Day10) following Drebrin inactivation in dendrites with >10% spine loss (45/123 dendrites) in dendrites with >10% spine loss (45/123 dendrites). Spine gain Day2 (baseline)=8.705±1.111%, n=32; Day3=5.871± 0.9229%, n=44 dendrites; Day10= 20.21±2.130%, n=30 dendrites. Day2 vs Day3 non significant, Day2 vs Day10 ***p < 0.001, Day3 vs Day10 ****p < 0.0001. Kruskal-Wallis test. e) Line graphs depicting a significant increase in the size of individuals dendritic spines 24 hours (Day3) and 1 week (Day10) following Drebrin inactivation in dendrites with >10% spine loss (45/123 dendrites). Spine sizes Day1=0.9457±0.024 (n=846 spines), Day2=1.00±0.023 (n=1025 spines), Day3=1.280±0.034 (n=934 spines); Day10=1.216±0.039 (n=631 spines). Day1 vs Day2 non-significant, Day2 vs Day3 ****p < 0.0001, Day2 vs Day10 ****p < 0.0001, Day3 vs Day10 non significant. Kruskal-Wallis test. f) Line graph depicting the averaged spine size for data showed in O). Note that spine loss mediated by Drebrin inactivation (orange bar) leads to the long-term enlargement of the remaining dendritic spines. ****p < 0.0001.

### Aßo-mediated spine elimination triggers a two-stage compensatory response

To examine whether these forms of compensation are relevant to AD, we first implemented an *in-vitro* model of β-amyloid-induced synapse loss [28] to track the emergence of spine compensation. Using confocal longitudinal imaging, we visualized dendritic regions before and after 3 and 5 hours of acute Aßo treatment in primary cultured neurons overexpressing Homer1c_GFP and tdTomato. We found Aßo triggered comparable levels of dendritic spine loss both at 3 and 5 hours (Fig. 3a.b). To investigate whether this was accompanied by the enlargement of the surviving dendritic spines, we measured the size of individual spines before and after Aßo application. Strikingly, we found a significant enlargement of remaining dendritic spines after 5 but not 3 hours of treatment (Fig. 3c.d) indicating that structural compensation does not occur concurrently with spine loss but emerges at a later stage. Similar to optogenetic spine elimination, we found that smaller spines were selectively enlarged following 5 hours of treatment with Aßo (Fig. 3e.f). Consistent with previous studies [33, 43], we found that although Aβo induced dendritic spine loss across the entire size distribution, small spines were slightly more susceptible at both 3 and 5 hours (Fig. 3g and Suppl. Fig. 3a).

**Figure 3.**
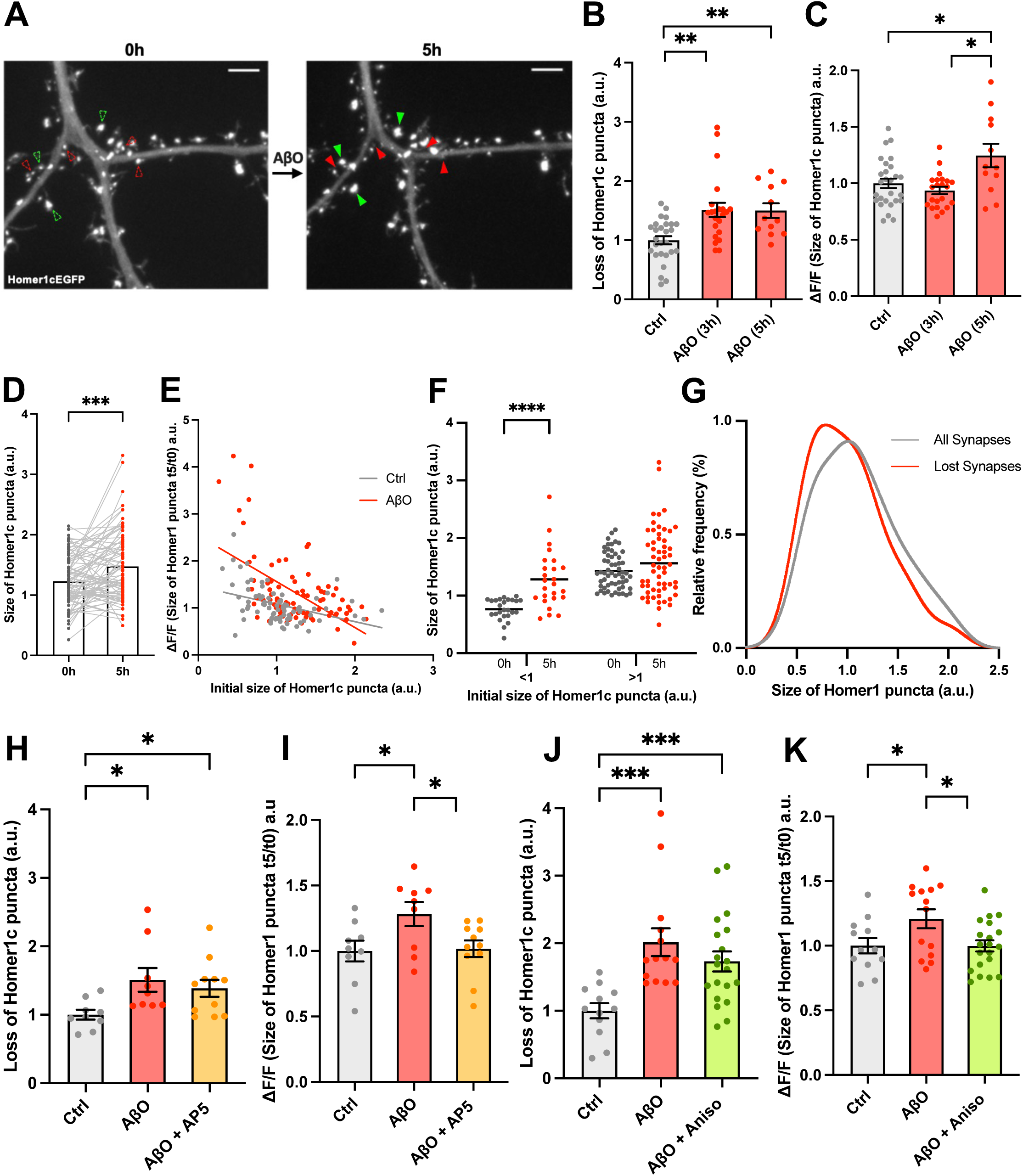
Mechanisms of spine compensation in an *in-vitro* Alzheimer’s disease model. a) Representative dendritic region of a primary cultured hippocampal neuron transfected with Homer1c-EGFP before (left) and after (right) 5 hours of Aβo treatment. Note that Aβo induces loss of dendritic spines (red arrowheads) and compensatory enlargement of dendritic spines in remaining synapses (green arrowheads) compared to baseline imaging (red and green hollow arrowheads). Scale bars 3µm. b) Bar graph showing a significant loss of Homer1c puncta at both 3 and 5 hours of Aβo treatment (3 hours: 19.82% ± 1.562 with a relative loss =1.511; 5 hours: 24.37% ± 2.015; relative loss =1.5) compared to the vehicle treated (DMSO) controls (13.11% ± 1.194 and 16.25% ± 1.836 at 3 and 5 hours, respectively; pooled data). **p <0.01, unpaired Kolmogorov-Smirnov test and unpaired t test with Welch’s correction respectively. Data points in bar graph corresponds to neurons. All data are ± SEM. c) Bar graphs representing the size of Homer1c puncta in surviving spines before (Ctrl; pooled for 3 and 5 hours) and after 3-hours (relative size 0.9366 ± 0.0323) or 5-hours (1.246 ± 0.1030) of Aβo treatment. respectively. *p < 0.05, unpaired t test with Welch’s correction. Data points in bar graph corresponds to neurons. All data are ± SEM. d) Bar graphs representing the increase in size of Homer1c puncta at the individual spine level before (1.230 ± 0.0454 a.u.) and after 5-hours (1.476 ± 0.06302 a.u.) of Aβo treatment respectively. ***p<0.001, paired Wilcoxon test. n = 83 spines from 12 neurons. e) Significant negative correlation observed between the initial size of Homer1c puncta at t=0 hours and its relative size after 5 hours in both Aβo-treated (r = - 0.4244) and vehicle controls (r = - 0.4393). ****p < 0.0001, Spearman’s correlation test. Simple linear regression analysis shows that slopes are significantly different (**p<0.01) between the two treatment groups. f) Graph showing that Homer1c puncta in smaller dendritic spines (brightness ratio < 1) are significantly enlarged after 5 hours, with no change for larger spines (brightness ratio >1). ****p < 0.0001, paired Wilcoxon test. Data points in bar graph corresponds to individual puncta. g) Frequency distribution for the size of lost Homer1c puncta compared to the size of all Homer1c puncta in the original population. Note that the distribution of the lost synapses is slightly shifted to the left indicating a preferential loss of smaller synapses. h) Bar graph showing a significant loss of Homer1c puncta in both Aβo only (positive control; 1.508 ± 0.1726, n=9) and Aβo + D-AP5 (1.385 ± 0.1249, n=11) treated neurons after 5 hours, relative to the vehicle (DMSO + D-AP5, n=9) treated controls. *p<0.05, unpaired t test with Welch’s correction. All data are ± SEM. i) Bar graphs showing that the relative size of Homer1c puncta in surviving spines was enlarged only in positive-control Aβo only treatment (1.282 ± 0.0916) but not in Aβo + D-AP5 (1.017 ± 0.0635) or vehicle controls. *p<0.05, unpaired t test with Welch’s correction. All data are ± SEM. j) Bar graph showing that the loss of Homer1c puncta was induced in both positive-control Aβo only (2.014 ± 0.2061, n=14) and Aβo + Anisomycin (1.730 ± 0.1485, n=20) treated neurons after 5 hours respectively relative to mean loss in vehicle-treated (DMSO + Anisomycin) controls (n=12). ***p <0.001, unpaired t test with Welch’s correction and unpaired Kolmogorov-Smirnov test respectively. All data are ± SEM. k) Bar graphs showing that the relative size of Homer1c puncta in surviving spines was significantly enlarged only in positive-control Aβo only treatment (1.208 ± 0.0729) but not in Aβo + Anisomycin (0.9986 ± 0.0434) or vehicle controls. *p < 0.05, unpaired t test with Welch’s correction. n values represent the number of neurons. All data are ± SEM.

To start investigating the mechanisms driving spine compensation, we took advantage of the time window between 3 and 5 hours of Aßo incubation where spine enlargement was likely taking place. By applying the pharmacological inhibitor AP5 (50μM) during the last two hours of the 5 hours-Aßo incubation, we found that the enlargement of dendritic spines depended on the activation of NMDA receptors (Fig. 3h.i). In addition, the compensatory enlargement of dendritic spines was also dependent on new protein synthesis (Fig. 3j.k). Likely due to the acute nature of the treatment, we failed to detect an increase in synaptogenesis at either 3 or 5 hours (Suppl. Fig. 4a.b), and assessment at 24 hours was precluded by reduced neuronal viability.

### In-vivo validation of Aßo-mediated spine elimination and compensation

Last, we investigated whether spine compensation occurs in *in-vivo* models of β-amyloid-induced synapse loss. As previously reported [1, 35, 43], we first verified that dendritic spine loss is accompanied by enlargement of remaining spines in the APP/PS1 model of AD. As dendritic spine loss only occurred in close proximity to amyloid plaques (<50 µm) [6, 17, 36], we explored whether enlargement of remaining spines is also restricted to this region. Using *in-vivo* two photon imaging to simultaneously image YFP-labelled dendrites and Methoxy-X04-labelled plaques in double transgenic APP/PS1xThy1-YFP mice (Fig. 4a), we confirmed that dendrites close to plaques not only have a reduction in spine density, but also show significantly enlarged spines (Fig. 4a-d). Because amyloid plaque deposition is stochastic, thus precluding the acquisition of baseline imaging prior to plaque deposition, it remained unclear whether the enlargement of the remaining spines originated from the loss of dendritic spines. To circumvent this issue, we longitudinally imaged the same dendritic region before and after the local injection of Aßo in the cortex of wild type C57BL/6 mice. We utilised an AAV-driven Cre-lox system (AAV9.hSyn.Cre, AAV9.flex.hSyn.EGFP-Homer1c and AAV9.CAG.flex_tdTomato) to sparsely express Homer1c and tdTomato - synaptic and structural markers, respectively. Following baseline imaging, we injected Aßo directly into the cortex and imaged the same dendritic region 24 hours and 1 week later (Fig. 4e and Suppl. Fig. 5a). As expected, we found that Aßo triggered loss of dendritic spines at both times points, though it only reached significance after 1 week (Fig. 4f.g). We then measured the size of individual spines before and after Aßo injection and found that retained dendritic spines were enlarged 24 hours and 1 week post-treatment, though again it only reached significance after 1 week (Fig. 4i.j). Critically, injection of vehicle control failed to induce dendritic spine loss or structural compensation (Fig. 4g-j). We then explored whether Aßo-mediated spine loss is followed by an enhancement in the formation of new dendritic spines. In agreement with the artificial system, we found that the loss of spines was followed by a significant increase in new spine formation at 1 week compared to vehicle control (Fig. 4h), suggesting activation of compensatory spinogenesis mechanisms to counteract dendritic spine loss.

**Figure 4.**
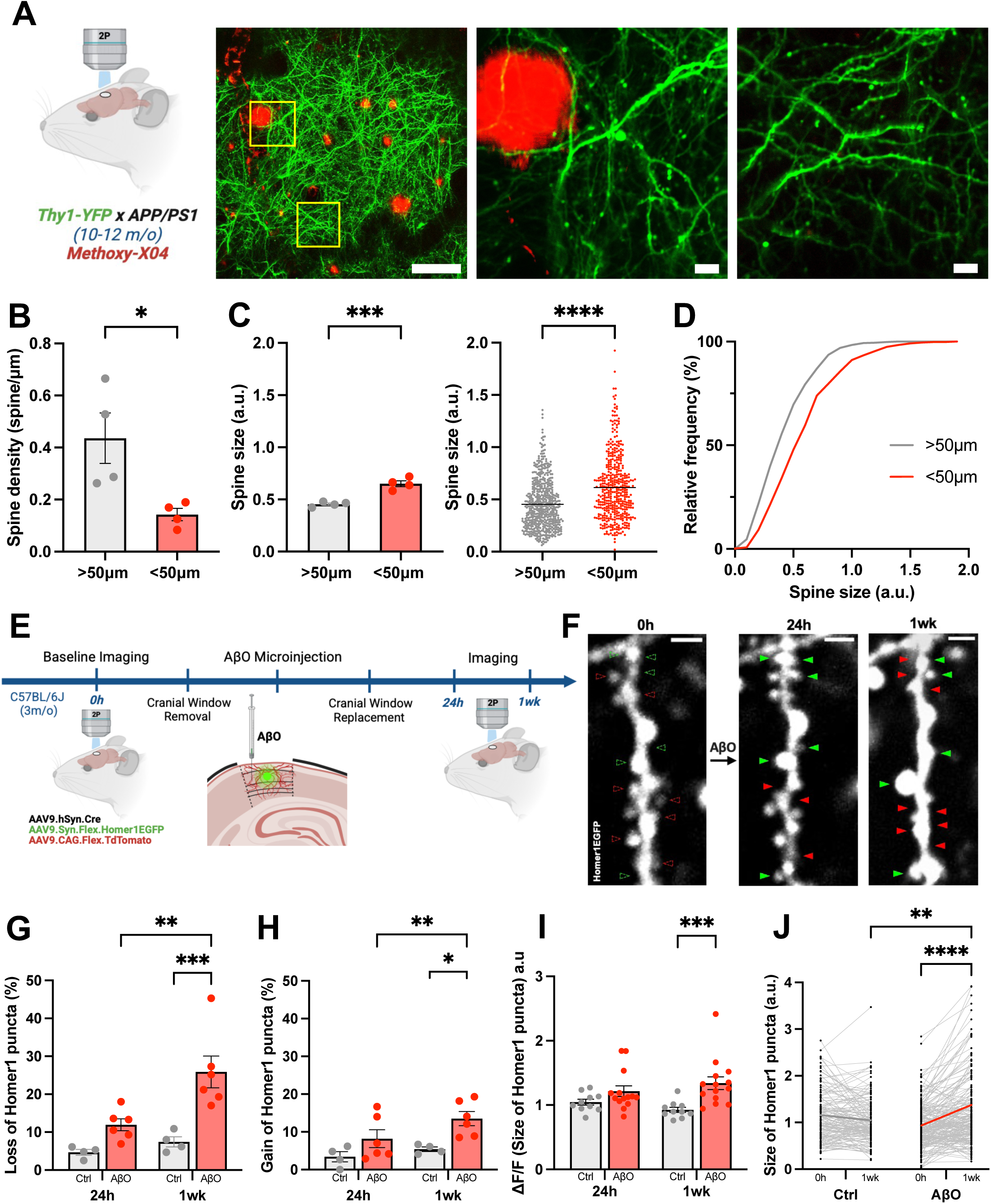
*In-vivo* dendritic spine compensation in Alzheimer’s disease models. a) Schematic *(left)* and images *(right)* showing the *in-vivo* two-photon imaging of YFP labelled dendrites in proximity of Methoxy-X04 labelled Aβ plaques in the APP/PS1xThy1-YFP double transgenic mice. Scale bars 100µm (*left* image) and 10µm (*middle* and *right* image). b) Bar graphs showing significant decrease in spine density within 50µm of Aβ plaques (0.1424 ± 0.02359, n= 4 mice) as compared to spines > 50µm (0.4358 ± 0.09705, n= 4 mice). Unpaired t-test, *p<0.05. All data are ± SEM. c) Graphs showing a significant enhancement in spine size within 50µm of Aβ plaques at the level of both the animal (*left*; <50µm= 0.6508 ± 0.02870, >50µm=0.4543 ± 0.01237, unpaired t-test, n = 4 mice, ***p<0.001) and individual dendritic spines (*right*; <50µm = 0.6148 ± 0.01473, n=438; >50µm = 0.4538 ± 0.00999, n=561; unpaired t-test, ****p<0.0001). Note that spine size analysis was conducted in same dendrites as in b). All data are ± SEM. d) Cumulative distribution of dendritic spines sizes in proximity (<50µm) or further away (>50µm) from Methoxy-X04 labelled Aβ plaques. e) Schematic displaying experimental workflow involving longitudinal *in-vivo* two-photon imaging before and after intra-cortical microinjection of oligomeric forms of the Amyloid beta peptide (Aßo) in anesthetised mice. f) *In-vivo* two-photon images of EGFP-Homer1c labelled dendrites in the cortex before and after intra-cortical Aßo injection. Note that Aßo induces dendritic spine loss (red arrowheads) and dendritic spines compensation – either enlargement of surviving spines or gain of new spines (green arrowheads) – compared to baseline imaging (red and green hollow arrowheads). Scale bars 3µm. g) Bar graphs showing higher loss of Homer1c puncta at 24 hours and 1 week postinjection of Aßo compared to DMSO vehicle controls (24hrs: Ctrl 4.675 ± 0.7728, Aßo 11.93 ± 1.589; 1 week: Ctrl 7.450 ± 1.333, Aßo 25.88 ± 4.178). Two-way ANOVA with Fishers LSD, n= 4/6 mice, 2-4 cells averaged per mouse, **p<0.01, ***p<0.001. All data are ± SEM. h) Bar graphs showing higher gain of Homer1c puncta at 24 hours and 1 week postinjection of Aßo compared to DMSO vehicle controls (24hrs: Ctrl 3.425 ± 1.349, Aßo 8.200 ± 2.351; 1 week: Ctrl 5.425 ± 0.5935, Aßo 13.52± 1.864). Two-way ANOVA with Fishers LSD, n= 4/6 mice, 2-4 cells averaged per mouse, *p<0.05, **p<0.01). All data are ± SEM. i) Relative size of Homer1 puncta demonstrates a significant increase at 1 week post-injection in Aßo treated dendrites compared to DMSO controls (24hrs: Ctrl 1.045 ± 0.04580, Aßo 1.223 ± 0.08846; 1 week: Ctrl 0.9255± 0.04098, Aßo 1.341± 0.09895). Two-way ANOVA with Fishers LSD, n= 10/14 dendrites, ***p<0.001. All data are ± SEM. j) Line graphs depicting a significant increase in the size of individuals dendritic spines in Aßo treated mice after 1 week, compared to DMSO controls (Ctrl: 24hrs 1.116 ± 0.04277, 1 week 1.032 ± 0.04050; Aßo: 24hrs 0.9270 ± 0.03360, 1 week 1.213 ± 0.05368). Two-way ANOVA with Tukey’s, n = 148/195 spines, **p<0.01, ****p<0.0001). All data are ± SEM.

## DISCUSSION

Here, we developed an optical tool for the selective elimination of dendritic spines to model this key feature of AD and identified a two-stage compensatory response: a rapid enlargement of surviving spines followed by delayed spine regeneration. More importantly, we showed that these structural adaptations also occur in AD models suggesting that neurons retain an intrinsic capacity to reverse early synaptic loss in AD, potentially contributing to cognitive resilience.

The integrity of dendritic spines plays a critical role in sustaining cognitive function in AD. Numerous studies have shown that spine loss strongly correlates with the severity of cognitive impairment in both patients and animal models [10, 30, 32, 40]. Conversely, the preservation of dendritic spines is associated with cognitive resilience — the ability to maintain normal cognition despite significant amyloid and tau pathology [2, 7, 44]. Despite the central importance of dendritic spines in AD, the neuronal mechanisms that compensate for their loss remain largely unknown [5]. Given that early synapse loss in AD is gradual and localized to dendrites in close proximity to plaques [6, 19, 36], it is unlikely to engage classical homeostatic plasticity mechanisms like synaptic upscaling, which require sustained and widespread reductions in neuronal activity [41]. Instead, local forms of structural compensation like the ones identified in this study are more likely to be implemented in the early stages of AD.

Although postmortem studies of AD patients and animal models have reported that dendritic spine loss is often accompanied by enlargement of the remaining spines [4, 5, 10, 33, 35, 43], this has typically been attributed to the preferential loss of small, vulnerable spines [4]. As such, the apparent increase in average spine size would simply reflect the surviving population of larger spines. However, by using longitudinal imaging of the same dendritic regions over time, we provide direct evidence that the spine enlargement observed in AD reflects a genuine compensatory response at the level of an individual spine. First, we established a causal link between spine loss—induced either by our artificial tool or by Aβo—and the compensatory enlargement of pre-existing spines at the single-spine level. Second, although Aβo preferentially targeted small spines, this alone could not account for the population-level increase in spine size. Notably, a 3-hour incubation with Aβo was sufficient to induce spine loss but did not result in population-wide spine enlargement. Interestingly, we observed that small spines were selectively enlarged—consistent with their higher plasticity [8, 21]—which further decreased their relative representation at the population level.

In addition to demonstrating that spine enlargement represents a *bona fide* form of structural compensation, we also provide the first mechanistic insight into this process. Specifically, we show that compensatory spine enlargement depends on NMDA receptor activation and de novo protein synthesis. While further studies are needed to fully characterize this form of compensation, the involvement of NMDA receptors suggests that active spines are preferentially engaged in this compensatory response, supporting its physiological relevance. This hypothesis is further supported by our findings showing that compensated spines show increased calcium responses. Moreover, the established role of NMDA receptors in driving calcium-dependent gene expression aligns with the requirement for new protein synthesis in this form of structural plasticity [11, 12].

Another potential mechanism of compensation for spine loss in AD is the *de novo* formation of dendritic spines. This regenerative response may represent a more robust form of structural compensation than the enlargement of existing spines, as it has the potential to restore overall spine density and re-establish lost synaptic connectivity [5]. Although previous studies using β-amyloidosis models of AD have provided indirect evidence of increased synaptogenesis [5, 18, 23], it remained unclear whether this represented a true compensatory response or was simply a consequence of APP overexpression, which is known to promote synaptogenesis during development [24, 45]. In this study, we provide direct evidence that both Aβo and optogenetically-induced spine loss drive de novo spine formation in wild-type animals, establishing a causal link between synapse loss and compensatory spinogenesis. While these findings support the idea that neurons retain the capacity to reverse early synaptic loss in AD, it remains to be determined whether the newly formed spines successfully re-establish functional synaptic connectivity.

A key question emerging from this study is whether the observed structural changes contribute to cognitive resilience—the capacity to maintain normal cognitive function despite substantial amyloid and tau pathology [37, 42]. While previous studies have shown that cognitively resilient individuals retain dendritic spine densities similar to age-matched controls [2, 7, 44], the mechanisms underlying this preservation remain unclear. Our findings suggest that an enhanced capacity for spinogenesis could play a role in counteracting spine loss in these individuals. Alternatively, resilience may arise from a higher baseline spine density or from reduced vulnerability of their spines to Aβo —for instance, due to the absence of critical Aβo-binding receptors. Notably, cognitive resilience has been associated with an increased proportion of thin, elongated spines [7, 44], which may represent newly formed spines actively engaging in synaptogenesis [14, 15]. This hypothesis is further supported by gene ontology analyses from multiple transcriptomic and proteomic studies of resilient individuals, which show enrichment in pathways related to cell-cell adhesion— processes known to be essential for synaptogenesis during development [3, 20, 27, 39].

In conclusion, we identified a two-stage compensatory response to dendritic spine loss: rapid enlargement of surviving spines followed by delayed spine regeneration. As these structural adaptations also occur in models of Alzheimer’s disease, our spine compensation framework offers a powerful platform to uncover fundamental molecular mechanisms that could be leveraged to reverse early synaptic loss and enhance cognitive resilience in AD.

## Acknowledgements

We would like to thanks Dr. Tomoaki Shirao for kindly providing the Drebrin1 plasmid, as well as Dr. Tobias Bonhoeffer who develop an earlier version of the optogenetic tool. This work was supported by the UK Dementia Research Institute (award number UK DRI-Edin0010] through UK DRI Ltd, principally funded by the Medical Research Council to P.O.; Dementia Australia Research Foundation Back Block Bards Project Grant to P.O.; and National Health and Medical Research Council Project Grant to P.O.

## Methods

### Animals

All experiments were performed under licenses approved by the UK Home Office according to the Animals (Scientific Procedures) Act and the University of Queensland Animal Ethics Committee. Female, adult Sprague-Dawley rats and their embryos (both, male and female at embryonic stage 18) were used for preparing the primary hippocampal neurons. Wistar rats postnatal day 4-5 were used for preparing organotypic hippocampal slice cultures. Male 12-week old CD1 (in-utero electroporated) and C57BL/6J (AAV-injected) mice were used for surgical procedures and in-vivo imaging experiments. Animals were typically housed in groups of 6 (2-3 following surgery), in a 12h/12h light dark cycle in standard cages containing appropriate enrichment and access to food and water.

### Primary neuronal cultures and transient cell transfection

Primary rat hippocampal neurons were prepared from Sprague-Dawley rat embryos (males and females, embryonic day 18). Briefly, isolated hippocampi were subjected to enzymatic digestion using 10U of papain suspension (Worthington Biochem. Co.) for 20min in a 37°C water bath and further mechanically dissociated by trituration using fire polished Pasteur pipettes. The single cell suspension was then plated at a density of 8 × 10^4^ cells per dish on Poly-L-lysine (0.5mg/mL; Sigma-Aldrich) coated 20mm glass bottom dishes (Cellvis) in plain Neurobasal plating medium (Gibco) supplemented with 5% Fetal Bovine Serum (FBS) (GE Healthcare), 1% L-glutamine (Gibco), 1% Penicillin-Streptomycin (Gibco) and 2 % B-27^TM^ supplement (Gibco) and incubated in a 37°C humidified tissue culture incubator with 5% CO_2_. Cultured neurons were fed twice a week with Neurobasal medium with no serum.

Cultured hippocampal neurons were transfected between days *in vitro* (DIV) 11 to 14 using Lipofectamine 2000 (Invitrogen) according to the manufacturer’s protocol and imaged DIV 15 onwards, after a minimum 48h post-transfection. Since neurons are post mitotic, exogenous plasmid DNA over expression was long lasting as well as non-toxic. The pCMV-Homer1c-EGFP plasmid was obtained from the Choquet lab, Bordeaux, France. Plasmid was purified using a DNA purification kit (Qiagen) prior to transfection.

### Organotypic slice culture preparation and biolistic transfection

Organotypic hippocampal slice cultures were prepared from postnatal day 3-5 Wistar rats. Hippocampal slices (400 μm thick) were prepared as described previously[38]. 25μg of Drebrin-KillerRed and 15 μg of eGFP were coated onto 12.5 mg of 1.6 μm gold beads. After 7 days *in vitro*, slice cultures were transfected by biolistic gene transfer (180psi) (Gene gun; Bio-Rad) as previously described[22], allowing for a minimum 72h post-transfection, before imaging.

### Aβ oligomer preparation and pharmacological treatments

Aβ oligomer was generated as previously[31]. Briefly, 1,1,1,3,3,3□hexafluoro□2□propanol (HFIP)□treated amyloid□β1□42 peptide (JPT Peptide Technologies GmbH, SP□Ab□07_0.5) was resuspended in fresh, sterile dimethyl sulfoxide (DMSO) (Sigma) as monomers, with sonication to facilitate resuspension. 0.5mg vial of the HFIP treated Aβ (1-42) peptide was dissolved in 22μl DMSO, aliquoted and stored at −80□C as a 5mM stock solution. For oligomeric Aβ treatment, the dissolved peptide stock was diluted in F-12 medium with L-glutamine (Gibco) to 100μM and then incubated overnight at 4°C. Following day, the soluble Aβ oligomers were used to treat primary cultured neurons at 2.5µM final concentration for either 3h or 5h at 37°C. For vehicle controls, DMSO was used instead of Aβ 1-42 peptide while keeping the rest of the protocol identical and processed parallelly. The concentration of Aβ oligomers is expressed in monomer equivalents. Oligomeric nature of the Aβ preparation was confirmed using Western blots with anti β-amyloid mouse antibody 6E10 (1:2,000 dilution; Covance).

Anisomycin (Tocris) and D-AP5 (Sigma-Aldrich) to inhibit local protein synthesis and NMDAR activity, were resuspended in fresh, sterile 100% DMSO and distilled water (UltraPureTM, Invitrogen) at stock concentration of 80mM and 20mM respectively, aliquoted and stored at −30□C. Both selective inhibitors were used at a working concentration of 50μM for the respective *in-vitro* experiments.

### In-utero electroporation

As previously described[29], time-mated CD1 pregnant dams were used at embryonic stage E15 (upper layer neurogenesis) for all experiments. Deep anaesthesia of mice for this recoverable procedure was induced with an intraperitoneal (IP) injection of Ketamine (100mg/mL, Parnell Laboratories)-Xylazine (20mg/mL, Troy Laboratories) mix at dosage of 120mg/kg and 10mg/kg respectively. Full anaesthesia was confirmed by toe pinch test, and the dams were placed on a heat pad. The eyes of the dams were covered with Vaseline (Unilever), in order to prevent drying. Before performing a laparotomy to expose the embryos from the abdominal cavity, the hair covering the abdomen was gently removed using a hair removal cream (Nair), the skin was sterilised with chlorohexidine (Pfizer), and sterile gauze (Medsure) was positioned on the skin, leaving a small circular window for the laparotomy. Once the embryos were exposed, each one of them was positioned so that the lateral telencephalic ventricles were visible through the uterine wall. Plasmid DNA with the addition of 0.0025% Fast Green dye (Sigma-Aldrich) was then microinjected with a Picospritzer II (Parker Hannifin) using a glass pulled pipette, into the right lateral ventricle. The tip of the glass pipette (Thin Wall Glass Capillaries 1.2mm OD / 0.90mm ID, WPI, FL), previously prepared using a Flaming/Brown micropipette puller (heat 495, pull 100, vel 100, Sutter Instrument Co., CA), was trimmed obliquely using forceps, before plasmid injection. The plasmids were then electroporated approximately into the right primary somatosensory cortex (S1) with 3mm-diameter microelectrodes (Nepagene) delivering five (100ms, 1Hz) approximately 36 V square wave pulses from an ECM 830 electroporator (BTX Harvard Apparatus). Once this procedure was completed for each embryo, the uterine horns were replaced inside the abdominal cavity and the incision was sutured closed twice. Animals were then subcutaneously injected with 1mL of sterile saline and recovered in a humidified chamber at approximately 28°C. For pain relief following electroporation, an edible buprenorphine solution (0.026mg/mL; Temgesic, Indivior) in a gel pack was positioned next to the recovering dam in the cage. The buprenorphine edible gel pack was prepared before the surgery by injecting sterile sucralose-water gel (MediGel®, ClearH2O®) with 0.2mL of edible buprenorphine solution. Dams were then monitored daily until they gave birth to live pups (at approximately E19/P0). The presence of fluorescent patches in S1 were checked using fluorescence goggles. Only the pups with fluorescent patch were kept till they reached desired experimental age and other pups without a fluorescence patch were excluded/sacrificed.

### Cranial Window Implantation

Following in utero electroporation, 12-week old male mice from a CD1 background were anaesthetised using either isoflurane (4% for induction and 1.7-2.5% for maintenance) or with an intraperitoneal injection of a sleep mix containing 0.05mg/mL Fentanyl (Sublimaze, Piramal), 5mg/mL Midazolam (Mylan), 1mg/mL Medetomidine (ilium) in sterile saline (Mini-Plasco®, B. Braun) at a dose of 10μl/g bodyweight before mounting them onto a stereotaxic frame (RWD Life Sciences). Body temperature was monitored and maintained at 37°C throughout the procedure. Mice were injected subcutaneously with Carprofen (20mg/kg) for pre-operative analgesia and eye ointment (Lubrithal, Dechra) was applied to prevent eyes drying out. The skin over the skull was disinfected with antimicrobial cleanser (Hibiscrub) and an incision was made in the skin which was then removed to expose the skull. A craniotomy (3-5mm in diameter) was made above the somatosensory cortex based on the stereotaxic coordinates −1.7mm (AP) and 2.5mm (ML) from Bregma, formed by drilling gently using a high-speed microdrill (RWD Life Sciences). The exposed brain was kept moist by applying sterile cortex buffer (7.3mg/mL NaCl, 0.37mg/mL KCl, 1.98mg/mL D-glucose, 2.38mg/mL HEPES, 0.29mg/mL CaCl2*2H2O, 0.49mg/mL MgSO4*7H2O in MilliQ H2O, adjusted to pH 7.4) before covering with a circular glass coverslip (5mm diameter). The window was sealed, and all the areas of exposed skull bone were covered using dental cement (Vertex Dental). A small custom-made titanium headplate was embedded in the acrylic for head-fixing the animal during imaging sessions.

### Viral Injection

For viral injection in 12-week old male mice from a C57BL/6J background, a craniotomy was first produced as described above, before a combination of adeno-associated viruses (AAV): AAV.hSyn.Cre.WPRE.hGH (Addgene_105553-AAV9, diluted 1:10,000); AAV-CAG-FLEX-tdTomato (RRID: Addgene_28306-AAV9, diluted 1:10) and AAV-FLEX-SYN1-EGFP-Homer1 (custom-made, VectorBuilder) were mixed in equal parts and injected into the somatosensory cortex (300nl) using a pulled glass micropipette and a microinjection syringe pump (RWD Life Sciences), at a rate of 100nl/min and a depth of ∼500μm below the dura. Sterile cortex buffer containing 1mg/ml Dexamethasone was applied to the exposed brain to reduce inflammation and a circular glass coverslip (5mm diameter) was placed over the craniotomy and fixed to the skull with a cyanoacrylate glue (Power Gel, Loctite). A custom-made steel headplate was implanted onto the exposed skull with glue and fixed with dental cement (Simplex, Kemdent). Subcutaneous injections of Buprenorphine (0.1mg/kg) and sterile saline were given for post-operative analgesia and rehydration. Mice were placed in a heat chamber to recover and returned to their cages for 3-4 weeks to allow for AAV expression and clearing of inflammation under the cranial window.

### Aβ oligomers intracortical microinjection

For Aβ oligomer microinjection mice were anaesthetised with an intraperitoneal injection of sleep mix containing 0.05mg/mL Fentanyl (Sublimaze, Piramal), 5mg/mL Midazolam (Mylan), 1mg/mL Medetomidine (ilium) in sterile saline (Mini-Plasco®, B. Braun) at a dose of 10μl/g bodyweight, and following baseline in-vivo two photon imaging were head-fixed under a surgical microscope. The cement around the cranial window was removed using a high-speed microdrill (RWD Life Sciences), before the glass coverslip was carefully lifted and removed. Mice were injected intracranially with 500nl of either Aβ oligomers (45ng total) or vehicle DMSO (0.4%) at a rate of 200nl/min into the cortex approximately 500μm (ML) from the edge of the two-photon imaging site. A new glass coverslip was then placed over the exposed brain and fixed using dental cement (Simplex, Kemdent). A subcutaneous injection of awake mix containing 0.4mg/mL Naloxone, 0.1mg/mL Flumazenil and 5mg/mL Atipamezole in sterile saline was given at a dose of 10ul/g bodyweight, along with sterile saline for recovery and rehydration. Mice were placed in a heat chamber to recover and returned to their cages. Subsequent two-photon imaging sessions were carried out at 24 hours- and 1 week-post injection.

### Microscopy

For *in-vitro* cultured hippocampal neurons, transfected live neurons were imaged between DIV 19 and DIV 24 in an Okolab stage-top incubator maintained at 37°C and 5% CO_2_. Before imaging, the culture medium was entirely replaced with artificial cerebrospinal fluid (ACSF; 145mM NaCl, 2mM CaCl_2_, 2mM MgCl_2_, 10mM HEPES, 10mM D-glucose and 5mM KCl in MilliQ H_2_O, adjusted to pH 7.4). Individual neurons were then imaged using a Nikon Plan Apochromat 100x/1.45 NA oil-immersion objective on an inverted spinning disk confocal microscope (Diskovery; Andor Technology, UK) built around a Nikon Ti-E body (Nikon Corporation, Japan) and equipped with two Zyla 4.2 sCMOS cameras (Andor Technology) and controlled by Nikon NIS software. Confocal laser at λ488nm (1.63mW at 10X) was used for excitation and imaging the Homer1c-eGFP. Secondary dendrites of co-transfected neurons were randomly imaged as z-stack with 0.4μm step size over a range of 5μm at sequential time points. A z-stack was captured before the Aβo or vehicle treatment (0h) and a second z-stack of the same dendritic region was captured at respective time-point (either 3h or 5h) following either treatment.

For *ex-vivo* organotypic hippocampal slice neurons, biolistically transfected neurons were longitudinally imaged between DIV 11 and 19 in ACSF (127mM NaCl, 2mM CaCl_2_, 1mM MgCl_2_, 25mM D-glucose, 2.5mM KCl, 25mM NaHCO_3_ and 1.25mM NaH_2_PO_4_*H_2_O) maintained at the temperature of 35°C and pH 7.4. Image stacks of transfected hippocampal pyramidal neurons were acquired using two-photon laser scanning microscopy (Ultima Multiphoton system, Bruker) equipped with a tunable Ti: Sapphire laser (Mai Tai or Chameleon Vision by Spectra Physics and Coherent, respectively) and Prairie View software (version 5.6). The image stacks (512 x 512 pixels; 0.68μm per pixel) with 0.5 μm z-steps were imaged using 60x immersion objective and an optical zoom of 10x, for each time-point. For Drebrin::KillerRed inactivation, the dendritic region of interest was irradiated with epifluorescence wide-field green light using a Mercury lamp (X-Cite® 120Q) until KillerRed photobleaching between 3-5 minutes.

For *in-vivo imaging*, apical dendritic stretches 10–100μm below the cortical surface, of co-labelled layer II/III pyramidal neurons were repeatedly imaged *in-vivo* in mice anesthetised with either isoflurane (4% for induction and 1.7-2.5% for maintenance) or with an intraperitoneal injection of a sleep mix (0.05mg/mL Fentanyl, 5mg/mL Midazolam and1mg/mL Medetomidine) at different time-points using two-photon laser scanning microscopy (Ultima Multiphoton system, Bruker) equipped with tunable Ti: Sapphire lasers and a x25 WI Nikon objective (NA 1.10; WD 2.0). For Drebrin::KillerRed inactivation, the dendritic region of interest was irradiated with epifluorescence wide-field green light using a Mercury lamp (X-Cite® 120Q) for 30 minutes.

### Image analysis

All the spine analyses were performed using FIJI (ImageJ). Unhealthy appearing neurons which displayed blebbing or loss of dendritic processes were excluded from analysis. Only the dendritic spines or Homer1c puncta appearing as clear protrusions adjacent to the dendrite, irrespective of their orientation relative to the imaging plane, were included in the analyses. Spine density was calculated as total number of spines or Homer1c puncta per measured dendritic length. The rate of spine loss and gain was calculated as percentages of spines that appeared and disappeared at the different time-points following treatments, relative to the corresponding number of spines at 0h. The integrated fluorescence intensity (brightness) of dendritic spines (either GFP or Homer1c puncta) on the z-stack maximal projections or the best focal section was used as a measure of spine size, as previously described [15]. The individual values for integrated fluorescence intensity in the spine or Homer1c puncta (within spines) were corrected for background and normalised to cytosolic, integrated fluorescence intensity in the adjacent dendritic shaft to account for differences in exogenous plasmid expression. Relative changes in dendritic spine or Homer1c puncta sizes were quantified as ratios of individual spine/punctum sizes following the respective manipulation to the initial size of the same spine prior to treatment. The individual values of relative change in size were then averaged per dendritic region for all treatment groups.

GCaMP analysis was based on work described previously[25]. Briefly, to measure spontaneous spine activity ROIs were drawn around spines of interest and mean gray value was measured simultaneously for all spines within the field of view for the duration of the experiment (5 minutes). ΔF/F was calculated by dividing the mean gray value over time for the spine (ΔF) with the averaged mean gray value of 5 ROIs drawn randomly along the dendritic shaft (F). This was done to ensure the observed changes are specific to spines and not the entire dendritic region.

### Statistics

All statistical tests were performed using Graph Pad Prism version 8 and above. Statistical tests were chosen based on tests for normality. The specific statistical analyses performed are listed in the respective figure legends. A p value of <0.05 was considered statistically significant. Depending on the experiments, either individuals’ spines (paired analysis) or neurons (independent samples) will correspond to the experimental unit. Correlation coefficients were calculated with a Spearman’s rank or Pearson’s correlation coefficient.

