## supplementary figures for "Direct Evidence for Dendritic Spine Compensation and Regeneration in Alzheimer’s Disease Models"

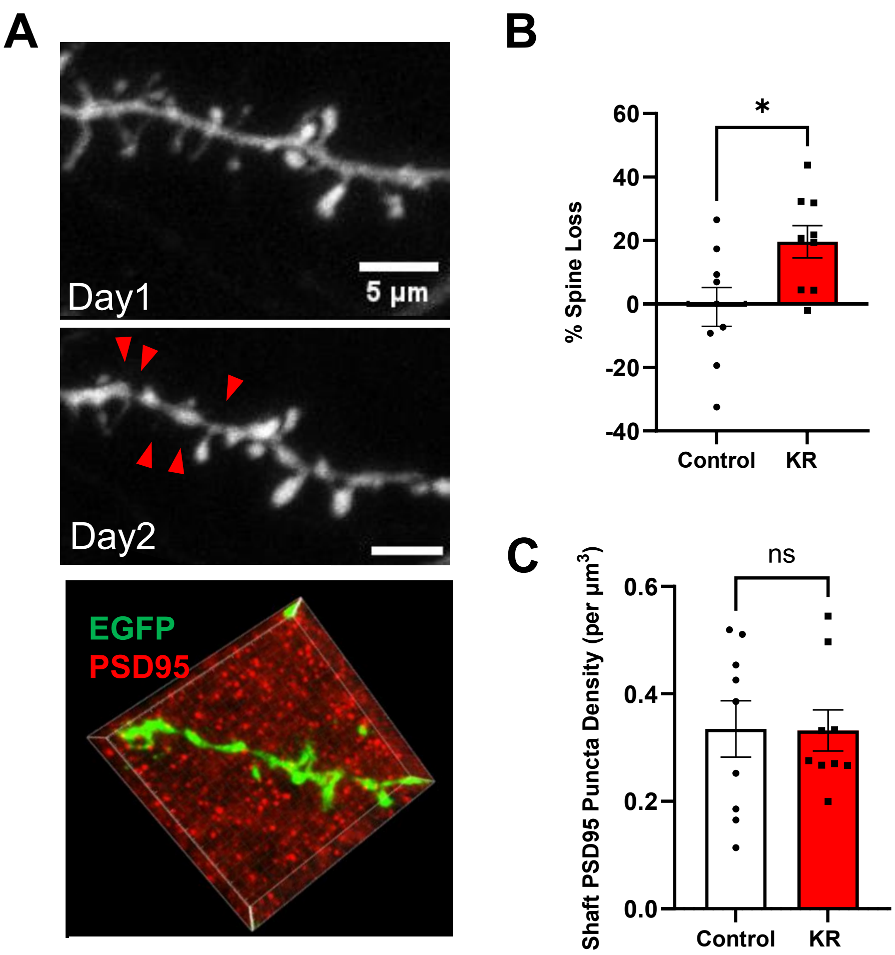


**Figure S1. Artificial elimination of dendritic spines does not lead to synapse relocation onto the shaft. A)** Two-photon images of hippocampal neuron in organotypic slice cultures overexpressing GFP and Drebrin::KillerRed before (*top*) and after (*middle*) the artificial elimination of spines (red arrowheads) and following post hoc immunohistochemistry for endogenous PSD95 (*bottom*). **B)** Bar graph showing a significant loss of dendritic spines 24hours post-Drebrin inactivation (% spine loss in controls (Ctrl)= -0.9041 ± 6.127%, n=9; % spine loss after Drebrin inactivation = 19.62± 5.049%. n=9 **p <0.05 unpaired t-test. **C)** Bar graph showing that spine elimination had no effect on the density of shaft PSD95 puncta suggesting that spine synapses did not relocate onto the shaft after spine elimination. Ctrl= 0.3347± 0.0523, n=9; Drebrin inactivation = 0.3318± 0.038. n=9 *p <0.05 unpaired t-test. All data are ± SEM. Note that a conversion from spine to shaft synapses would have led to an increase in PSD95 puncta in dendritic shaft.

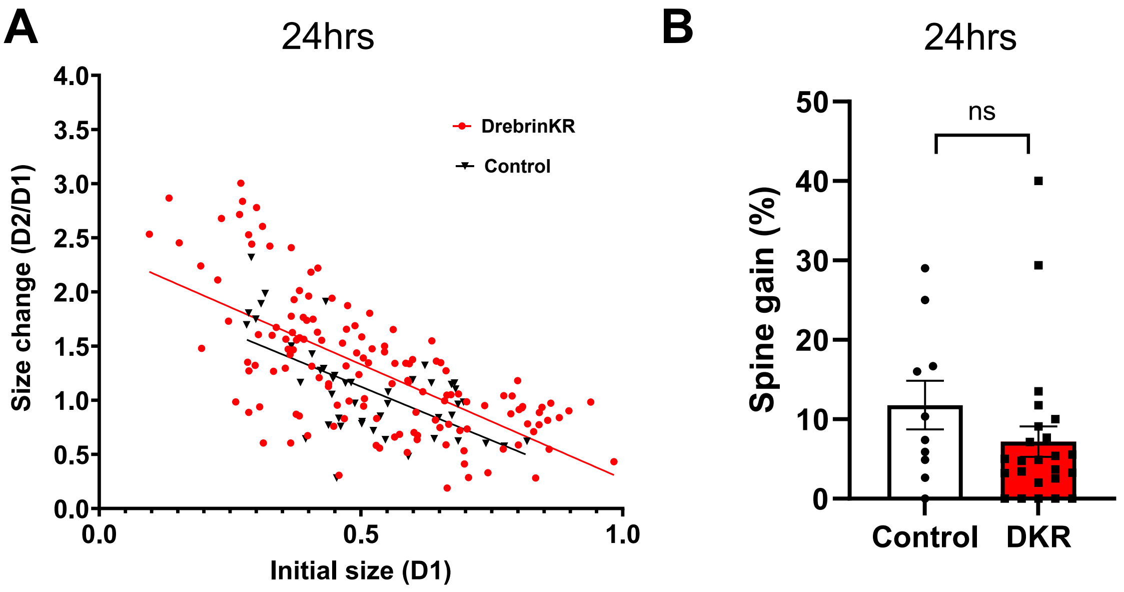


**Figure S2**. **Optogenetic spine elimination leads to enlargement of small spines but has no effect on spine gain 24 hours post-treatment**. **A)** Significant negative correlation observed between the initial spine size at Day 1 and spine compensation at Day 2 for controls (r squared =0.3976) and following Drebrin inactivation (r squared =0.444). Simple linear regression analysis shows that slopes from Ctrl and DKR significantly deviate from zero (***p<0.01). **B)** Bar graph showing no effect on dendritic spine gain 24hours post-Drebrin inactivation. Spine gain in controls (Ctrl)= 11.78± 3.055 %, n=10; spine gain after Drebrin inactivation = 7.185± 1.917%. n=24. p =0.2051 unpaired t-test. All data are ± SEM.

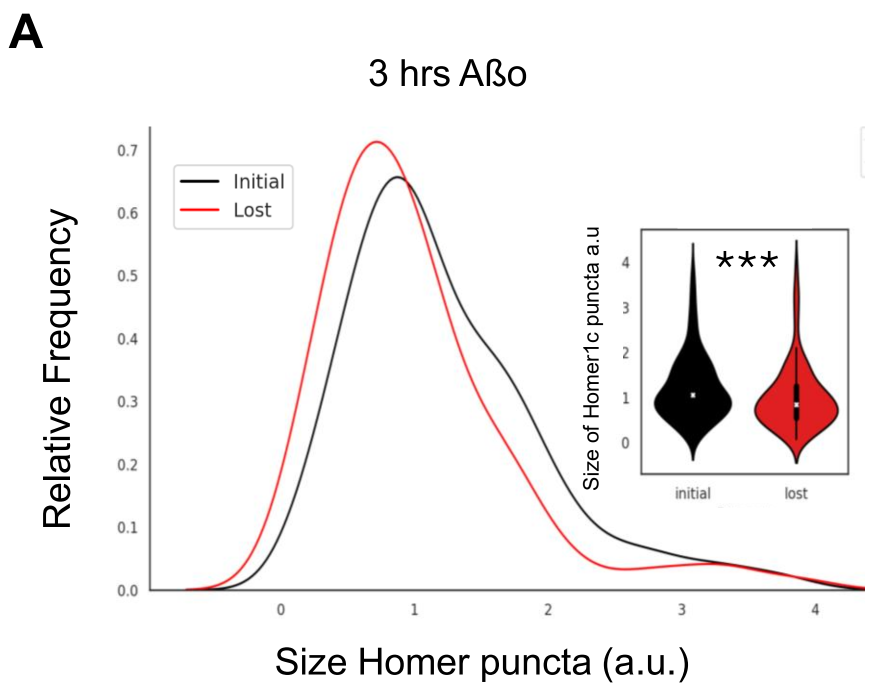


**Figure S3. Smaller dendritic spines are more vulnerable to Aβo treatment in primary hippocampal neuron cultures. A)** Frequency distribution for the size of lost Homer1c puncta compared to the size of all Homer1c puncta in the original population. Note that the distribution of the lost synapses is slightly shifted to the left indicating a preferential and significant loss of smaller synapses (inset). *** $p<0.001$.

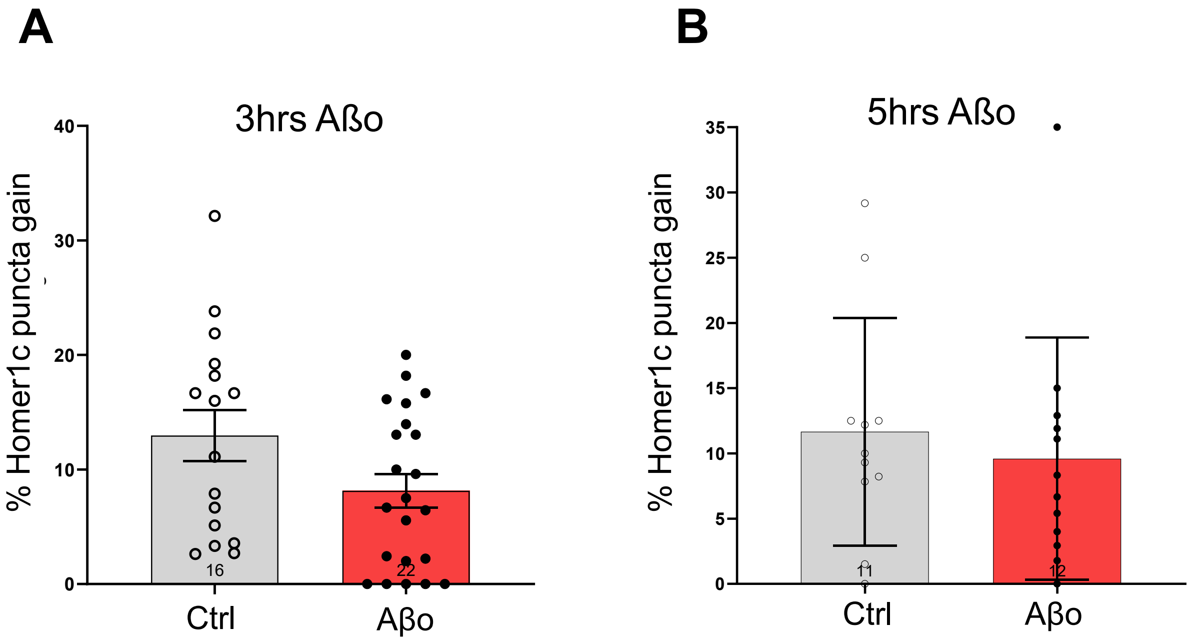


**Figure S4 No difference in gain of Homer1c puncta within dendritic spines following Aβo treatment in primary hippocampal neuron cultures.** No significant difference in gain of Homer1c puncta after both **(A)** 3-hours (8.148 ± 1.451) and (**B**) 5-hours (9.588 ± 2.682) of Aβo treatment compared to the vehicle controls (12.98 ± 2.228 and 11.66 ± 2.628), respectively. All data are ± SEM.

A

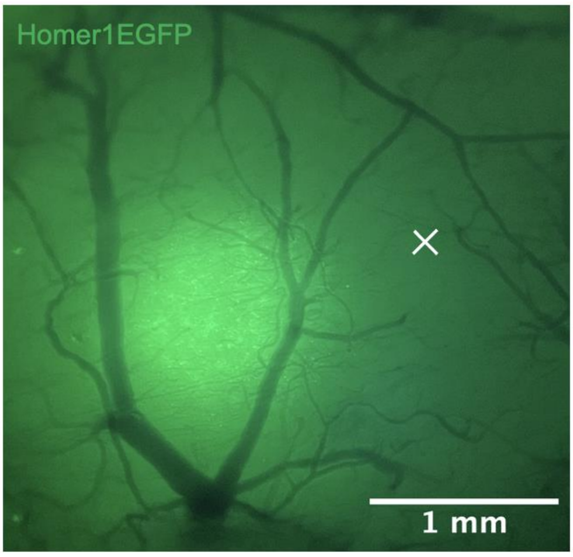


**Figure S5. Aβo injection sites relative to HomerEFGP-expressing neurons**. **(A)** Epifluorescent image following cranial window removal of HomerEGFP two-photon imaging site in the somatosensory cortex 4 weeks-post surgery, used to guide Aβo injection (white X).
